# Evaluating sepsis definitions for clinical decision support against a definition for epidemiological disease surveillance

**DOI:** 10.1101/648907

**Authors:** Anthony L. Lin, Mark Sendak, Armando D. Bedoya, Meredith E. Clement, Nathan Brajer, Joseph Futoma, Hayden B. Bosworth, Katherine A. Heller, Cara L. O’Brien

## Abstract

Epidemiological definitions for sepsis have been used to improve disease surveillance and assist in national assessment of sepsis detection and treatment. However, epidemiological definitions are often optimized for retrospective review and alignment of epidemiological and real-time clinical decision support definitions is unknown. This 15-month retrospective observational study involving 43,046 adult patient encounters and 30,973 unique patients compared cohorts captured by the EHR-based Sepsis-1, Sepsis-3, and Duke Adult Sepsis definitions. We assessed hospital length of stay, in-hospital mortality, admission to the intensive care unit, and sensitivity and specificity of these definitions compared to the Centers for Disease Control and Prevention epidemiological ‘gold-standard’ definition. Both Duke Adult Sepsis and Sepsis-3 maintained a high specificity, but the former had higher sensitivity. By directly comparing definitions designed for clinical decision support with an epidemiological disease surveillance tool, this work provides guidance to health systems seeking to implement clinical decision support to improve sepsis management.

## INTRODUCTION

Sepsis is a leading cause of morbidity and mortality in the United States, affecting 1.7 million adults and contributing to 250,000 deaths annually.^1^ Due to the lack of a clear diagnostic marker for sepsis, diagnosis has relied predominantly on clinical judgement based on suspicion for infection and organ dysfunction. In 1991, the American College of Chest Physicians/Society of Critical Care Medicine released a consensus definition for sepsis that relied on the Systemic Inflammatory Response Syndrome (SIRS) criteria.^2^ That definition was revised in 2016 with the Third Internal Consensus Definitions for Sepsis and Septic Shock (Sepsis-3) to reflect a growing understanding of the pathobiology of sepsis and delineate the difference between an uncomplicated infection and one resulting in end-organ dysfunction.^3^ The taskforce also released a new risk stratification tool to improve detection. The quick Sepsis-related Organ Failure Assessment (qSOFA) was shown to be more accurate than SIRS in predicting adverse clinical outcomes.^4^

In 2017, a Centers for Disease Control and Prevention (CDC)-funded consortium published a study that used a new definition for sepsis optimized for surveillance that relied on objective clinical data elements abstracted from electronic health records (EHRs).^1^ The CDC definition had greater predictive validity in estimating sepsis burden than the use of administrative codes as it was unaffected by underlying biases that may influence coding practices, such as increasing sepsis awareness and financial incentives.^5^

The CDC definition uses a blood culture order as an initial criterion. Case identification is predicated on four consecutive days of antibiotic administration and signs of organ dysfunction occurring within ±2 days of that blood culture order. As a result, identifying sepsis using the CDC definition is optimized for retrospective disease surveillance and cannot be used to identify cases in real-time. A real-time definition optimized for clinical decision support (CDS) can prompt meaningful clinical action at the moment that criteria for the definition are met. The CDC definition identifies patients up to 4 days after the time at which clinical action is needed. Health systems hoping to employ CDS to improve sepsis detection and treatment must align criteria that drive CDS with validated epidemiology measures such as the CDC definition.

The current study involves two analyses. First, cohorts identified by sepsis definitions that can be implemented in real-time are compared across a variety of clinical outcomes. Second, these cohorts are compared to patients identified by the epidemiological CDC definition. The study aims to provide guidance to health systems hoping to implement CDS to improve sepsis management.

## METHODS

This was a retrospective observational study of all adult (age ≥18 years) inpatient admissions at a large tertiary-care academic hospital with over 900 inpatient beds. The study was approved by our institutional review board (IRB Pro00080914). Data was obtained over a 15-month period, from October 1, 2014 to December 31, 2015. For these encounters, demographic information, comorbidities, vital sign measurements, laboratory results, medication administrations, and admission/discharge/transfer times were extracted from the EHR reporting database Clarity (Epic Systems) using Structured Query Language.

### Definition computation and statistical analysis

We used the Hospital Toolkit for Adult Sepsis Surveillance guide to compute cases of CDC Adult Sepsis Events.^6^ The Sepsis-1 and Sepsis-3 definitions were computed using previously-validated SIRS^7^ and qSOFA^4^ criteria. An order for any culture served as a proxy for clinician suspicion for infection to enable the Sepsis-1 and Sepsis-3 definition to be automatically computed from the EHR without manual chart review (see e-Table 1 for further details on implementation).

**Table 1.** Case identification and patient outcomes based on different sepsis definition criteria. CDC = Centers for Disease Control and Prevention, IQR = interquartile range, ICU = intensive care unit.

|  | Encounters, n (%) | Hospital length of stay (d), median (IQR) | In-hospital mortality, n (%) | ICU admission, n (%) |
| --- | --- | --- | --- | --- |
| **Sepsis-1** | 13,358 (31) | 6.9 (3.9-12.8) | 1,031 (8) | 3,989 (30) |
| **Sepsis-3** | 7,110 (17) | 8.3 (4.5-16.3) | 947 (13) | 3,201 (45) |
| **Duke Adult Sepsis** | 9,184 (21) | 7.3 (4.1-14.6) | 936 (10) | 3,167 (35) |
| **CDC Adult Sepsis Events** | 3,308 (8) | 13.1 (7.3-25.6) | 493 (16) | 1,789 (54) |
| **All Encounters** | 43,046 (100) | 4.0 (2.4-7.0) | 1,276 (3) | 8,132 (19) |

A new definition was also proposed by clinical experts at our local institution. Like other institutions,^8^ this resulted from a need to identify a real-time definition that accurately captured patients who could benefit from standard protocolized sepsis care bundles.^9^ The definition, termed [name deleted to maintain integrity of review process] Adult Sepsis, is conceptually similar to the Centers for Medicare and Medicaid Services (CMS) definition used to calculate timely delivery of those care bundles. The definition is met when ≥2 SIRS criteria are present, a blood culture is ordered (regardless of result), and ≥1 of the following elements of end-organ dysfunction is present: (1) creatinine >2.0 mg/dL, (2) international normalized ratio >1.5, (3) total bilirubin >2.0 mg/mL, (4) platelets <100 x 10^9^/L, (5) lactate ≥2 mmol/L, or (6) systolic blood pressure <90 mmHg or decrease in systolic blood pressure by >40 mmHg (see e-Table 1 for further details). Similar to the EHR-based Sepsis-1 and Sepsis-3 definitions, Duke Adult Sepsis is designed for real-time implementation within clinical decision support.

The cohort size, clinical characteristics, and setting of presentation (grouped into emergency department, inpatient wards, and intensive care unit) of each definition cohort were compared using aggregate statistics. Outcomes measured included hospital length of stay (LOS), in-hospital mortality, and rate of admission to the intensive care unit (ICU). Additionally, sensitivity, specificity, positive predictive value (PPV) and negative predictive value (NPV) were calculated for each real-time definition using CDC Adult Sepsis Events as the gold standard. A two-sided t test (α=0.05) was used to assess for statistical significance between cohorts. All sepsis definitions and statistical analyses were completed using R (version 3.4.1, 2017).

## RESULTS

In total, 43,046 unique inpatient encounters among 30,973 unique patients were captured (Table 1). A detailed overview of demographics, clinical characteristics, and 1-year comorbidities can be found in e-Table 2. The Sepsis-1 cohort was the largest (n=13,358, 31%), followed by the Duke Adult Sepsis cohort (n=9,184, 21%). The CDC Adult Sepsis Events cohort was the most restrictive, capturing only 8% (n=3,308) of total unique encounters. All four sepsis cohorts had greater hospital LOS, in-hospital mortality rates, and ICU admission rates than the average inpatient encounter. Of the four cohorts, Sepsis-1 captured patient encounters with the lowest hospital LOS, mortality rate, and ICU admission rate (6.9 days, 8%, and 30%, respectively). The CDC Adult Sepsis Events cohort had the highest hospital LOS, mortality rate, and ICU admission rate (13.1 days, 16%, and 54%, respectively). In e-Table 3, we report the distribution by setting of presentation for each definition at our local institution.

**Table 2.** Performance of the 3 real-time EHR-based definitions in detecting CDC Adult Sepsis Events. CDC = Centers for Disease Control and Prevention, CI = confidence interval, NPV = negative predictive value, PPV = positive predictive value.

| Definition |  | Positive CDC Adult Sepsis Event | Negative CDC Adult Sepsis Event | Combo Values (95% CI) |
| --- | --- | --- | --- | --- |
| **Sepsis-1** | Positive<br>Negative | 2,140<br>1,168 | 10,173<br>29,565 | Sensitivity: 0.647 (0.630-0.663)<br>Specificity: 0.744 (0.740-0.748)<br>PPV: 0.174 (0.167-0.181)<br>NPV: 0.962 (0.960-0.964) |
| **Sepsis-3** | Positive<br>Negative | 1,236<br>2,072 | 4,672<br>35,066 | Sensitivity: 0.374 (0.357-0.390)<br>Specificity: 0.882 (0.879-0.886)<br>PPV: 0.209 (0.199-0.220)<br>NPV: 0.944 (0.942-0.947) |
| **Duke Adult Sepsis** | Positive<br>Negative | 1,938<br>1,370 | 6,033<br>33,705 | Sensitivity: 0.586 (0.569-0.603)<br>Specificity: 0.848 (0.845-0.852)<br>PPV: 0.243 (0.234-0.253)<br>NPV: 0.961 (0.959-0.963) |

Table 2 illustrates how well Sepsis-1, Sepsis-3, and Duke Adult Sepsis identified cases confirmed by CDC Adult Sepsis Events criteria. The goal was to assess each definition’s ability to identify cases in real-time that ultimately met the validated epidemiology measure. Sensitivities for the three EHR-based definitions ranged from 0.374 (Sepsis-3) to 0.647 (Sepsis-1) while specificities ranged from 0.744 (Sepsis-1) to 0.882 (Sepsis-3). PPV was highest for the Duke Adult Sepsis definition (0.243) while NPV was similar between all 3 definitions, ranging from 0.944 (Sepsis-3) to 0.962 (Duke Adult Sepsis).

## DISCUSSION

Our findings emphasize the differences in patient population and clinical outcomes between sepsis definitions, which is well-described in existing literature.^10–12^ We also demonstrate the difficulty aligning sepsis treatment programs that must identify patients in real-time with sepsis disease surveillance, which requires information from the entire hospital course. Sepsis-3 had the lowest sensitivity of the 3 definitions and identified the smallest proportion of cases in the emergency department and the greatest proportion of cases in the ICU. These findings support previous work that criticize the definition for capturing sepsis late in its clinical course.^13^ As such, the definition captures the sickest patients; in this study these patients had the highest in-hospital mortality and ICU admission rates. Between Sepsis-1 and Duke Adult Sepsis, Sepsis-1 had the higher sensitivity while Duke Adult Sepsis had the higher specificity and PPV.

Local stakeholders were interested in identifying the optimal sepsis criteria for CDS to facilitate treatment in a timely fashion. The Duke Adult Sepsis definition meets these criteria by being computable within the EHR and balancing sensitivity and specificity using the CDC definition as a benchmark. The Duke Adult Sepsis definition maintains high specificity and sensitivity compared to Sepsis-1 and Sepsis-3.

Our work has several limitations. First, it is a single-center investigation. Though our inclusion criteria were kept intentionally broad to be most representative of the general hospital population, our findings may not generalize to other study settings.^8^ Second, as we sought to identify a sepsis definition that could be automatically computed from discrete elements in the EHR, we did not explore unstructured data (such as text from notes or reports). As technology behind natural language processing improves, future work could investigate the benefit of incorporating unstructured text in sepsis detection.

As advised by the 2016 sepsis definition framework,^10,14^ it is important to understand the tradeoffs of various sepsis definitions. Prior studies evaluated competing CDS definitions, whereas the current study is the first to compare CDS definitions to an epidemiology surveillance definition. Our findings can inform sepsis management and quality improvement programs at sites hoping to align CDS with epidemiology surveillance. Future work will test the Duke Adult Sepsis definition prospectively at our local institution. Ultimately, further research will be required to generalize this approach to other institutions. By sharing our analyses at our local institution, we hope to support other institutions interested in better aligning their sepsis CDS with current epidemiological measures.

