## Supplemental Tables for "Evaluating sepsis definitions for clinical decision support against a definition for epidemiological disease surveillance"

**e-Table 1.** The clinical criteria and time frames used to calculate each sepsis definition. CDC = Centers for Disease Control and Prevention, SIRS = Systemic Inflammatory Response Syndrome, qSOFA = quick Sepsis-related Organ Failure Assessment.

| | |
|---|---|
| Sepsis-1 | 2 or more SIRS criteria:<br>• Temperature >38.0°C or <36.0°C (6 hours)<br>• Heart rate >90 beats/min (6 hours)<br>• Respiratory rate >20 breaths/min (6 hours)<br>• White blood cell count >12,000/$mm^3$, <4,000/$mm^3$, or >10% immature neutrophils (24 hours)<br>Plus, any culture ordered:<br>• Blood, urine, sputum, or cerebrospinal fluid culture ordered (24 hours)<br>Plus, 1 element of end organ damage:<br>• Creatinine >2.0 mg/dL (24 hours)<br>• INR >1.5 (24 hours)<br>• Total bilirubin >2.0 mg/mL (24 hours)<br>• Platelets <100 x $10^9$/L (24 hours)<br>• Lactate ≥2 mmol/L (24 hours)<br>• Systolic blood pressure <90 mmHg or decrease in systolic blood pressure by >40 mmHg (6 hours) |
| Sepsis-3 | 2 or more qSOFA criteria:<br>• Systolic blood pressure ≤100 mmHg (6 hours)<br>• Respiratory rate ≥22 breaths/min (6 hours)<br>• Alert Voice Pain Unresponsive scale other than “Alert” (6 hours)<br>Plus, any culture ordered:<br>• Blood, urine, sputum, or cerebrospinal fluid culture ordered (24 hours) |
| Duke Adult Sepsis | 2 or more SIRS criteria:<br>• Temperature >38.0°C or <36.0°C (6 hours)<br>• Heart rate >90 beats/min (6 hours)<br>• Respiratory rate >20 breaths/min (6 hours)<br>• White blood cell count >12,000/$mm^3$, <4,000/$mm^3$, or >10% immature neutrophils (24 hours)<br>Plus, blood culture ordered:<br>• Blood culture ordered (24 hours)<br>Plus, 1 element of end organ damage:<br>• Creatinine >2.0 mg/dL (24 hours)<br>• INR >1.5 (24 hours)<br>• Total bilirubin >2.0 mg/mL (24 hours)<br>• Platelets <100 x $10^9$/L (24 hours)<br>• Lactate ≥2 mmol/L (24 hours) |

| | |
|---|---|
| | • Systolic blood pressure <90 mmHg or decrease in systolic blood pressure by >40 mmHg (6 hours) |
| CDC Adult Sepsis Events | Blood culture obtained<br><br>Plus, new antibiotic starting within ±2 days of blood culture day, followed by a total of ≥4 consecutive antibiotic days<br><br>Plus, CDC-specified organ dysfunction (±2 days of blood culture order day):<br>• Initiation of a new vasopressor infusion<br>• Initiation of invasive mechanical ventilation<br>• Serum creatinine ≥200% of encounter baseline<br>• Total bilirubin ≥2 mg/dL AND ≥150% of encounter baseline<br>• Platelet count <100 cells/uL AND <50% of encounter baseline (baseline must be ≥100 cells/uL)<br>• Serum lactate ≥2 mg/dL |

**e-Table 2.** Detailed clinical and demographic characteristics of the total study population and each definition cohort. Comorbidities are grouped according to the Agency for Healthcare Research and Quality Elixhauser Comorbidity Software (version 3.7, 2017) based on International Classification of Diseases (ICD)-9 and ICD-10 codes. CDC = Centers for Disease Control and Prevention, SD = standard deviation.

| | **All Encounters** | **Sepsis-1** | **Sepsis-3** | **Duke Adult Sepsis** | **CDC Adult Sepsis Events** |
|---|---|---|---|---|---|
| **Encounters, n (%)** | 43,046 (100) | 13,358 (31) | 7,110 (17) | 9,184 (21) | 3,308 (8) |
| **Age (yr), mean±SD** | 55.5±18.7 | 58.4±17.5 | 59.2±16.9 | 58.5±17.2 | 57.6±16.7 |
| **Female, n (%)** | 22,672 (52) | 8,873 (47) | 5,265 (46) | 6,336 (45) | 2,626 (42) |
| **Race, n (%)** | | | | | |
| White | 26,920 (63) | 8,440 (63) | 4,584 (64) | 5,713 (62) | 1,961 (62) |
| Black | 12,634 (30) | 4,135 (31) | 2,077 (29) | 2,914 (32) | 972 (31) |
| Missing/Other | 3,492 (8) | 783 (6) | 449 (6) | 557 (6) | 218 (7) |
| **Comorbidities, n (%)** | | | | | |
| Congestive Heart Failure | 6,324 (15) | 2,689 (20) | 1,725 (24) | 1,997 (22) | 784 (24) |
| Valvular Disease | 5,860 (14) | 2,338 (17) | 1,486 (21) | 1,619 (18) | 627 (19) |
| Peripheral Vascular Disease | 4,778 (11) | 1,842 (14) | 1,054 (15) | 1,292 (14) | 475 (14) |
| Hypertension | 20,345 (47) | 7,367 (55) | 3,787 (53) | 5,156 (56) | 1,807 (55) |
| Neurological Disorders | 7,485 (17) | 3,308 (25) | 2,064 (29) | 2,536 (28) | 1,037 (31) |
| Pulmonary Circulation Disorders | 7,728 (18) | 3,105 (23) | 1,603 (23) | 2,235 (24) | 747 (23) |
| Diabetes Mellitus | 9,825 (23) | 3,868 (29) | 1,984 (28) | 2,817 (31) | 1,030 (31) |
| Renal Failure | 6,885 (16) | 3,147 (24) | 1,621 (23) | 2,352 (26) | 820 (25) |
| Solid Tumor without Metastasis | 5,252 (12) | 1,571 (12) | 833 (12) | 1,019 (11) | 369 (11) |
| Coagulopathy | 5,020 (12) | 2,479 (19) | 1,421 (20) | 1,970 (21) | 882 (27) |
| Obesity | 6,162 (14) | 2,056 (15) | 1,043 (15) | 1,465 (16) | 524 (16) |
| HIV/AIDS | 444 (1) | 201 (2) | 86 (1) | 162 (2) | 43 (1) |
| Fluid and Electrolyte Disorders | 11,353 (26) | 5,290 (40) | 2,904 (41) | 4,047 (44) | 1,627 (49) |
| Anemia | 10,326 (24) | 4,449 (33) | 2,282 (32) | 3,378 (37) | 1,280 (39) |
| Depression | 7,045 (16) | 2593 (19) | 1,330 (19) | 1,871 (20) | 683 (21) |
| **Admission Source, n (%)** | | | | | |
| Self-Admit | 33,472 (78) | 9,522 (71) | 4,729 (67) | 6,395 (70) | 2,016 (64) |
| Transfer from Outside Hospital | 5,479 (13) | 2,502 (19) | 1,673 (24) | 1,847 (20) | 815 (26) |
| Healthcare Facility | 769 (2) | 396 (3) | 243 (3) | 335 (4) | 128 (4) |
| Outpatient Clinic | 2,938 (7) | 914 (7) | 453 (6) | 597 (7) | 188 (6) |
| Missing/Other | 388 (1) | 24 (<1) | 12 (2) | 10 (<1) | 4 (<1) |
| **Admission Type, n (%)** | | | | | |
| Routine | 13,210 (31) | 1,949 (15) | 1,188 (17) | 784 (9) | 313 (10) |
| Urgent | 11,548 (27) | 3,616 (27) | 1,954 (27) | 2,403 (26) | 879 (28) |
| Emergent | 18,288 (42) | 7,793 (58) | 3,968 (56) | 5,997 (65) | 1,959 (62) |
| **Discharge Destination, n (%)** | | | | | |
| Home | 35,625 (83) | 9,137 (68) | 4,176 (59) | 5,896 (64) | 1,675 (53) |
| Transfer to Outside Hospital | 221 (1) | 102 (1) | 82 (1) | 80 (1) | 36 (1) |
| Healthcare Facility | 4,819 (11) | 2,493 (19) | 1,553 (22) | 1,804 (20) | 770(24) |
| Deceased | 1,276 (3) | 1,031 (8) | 947 (13) | 936 (10) | 493 (16) |
| Missing/Other | 1,105 (3) | 595 (4) | 352 (5) | 468 (5) | 177 (6) |

**e-Table 3.** The distribution of each definition across Emergency Department (ED), Intermediate Wards, and Intensive Care Units (ICU) as well as time from setting presentation to meeting the clinical criteria of each definition. CDC = Centers for Disease Control and Prevention, IQR = interquartile range.

| | **Encounters, n (%)** | **Average # of encounters per day, n** | **Time from setting presentation to meeting criteria (h), median (IQR)** |
|---|---|---|---|
| **Sepsis-1** | | | |
| ED | 4,188 (31) | 9.2 | 1.8 (0.9-3.6) |
| Intermediate Wards | 6,904 (52) | 15.1 | 11.6 (3.2-38.7) |
| ICU | 2,266 (17) | 5.0 | 6.1 (1.9-21.6) |
| **Sepsis-3** | | | |
| ED | 894 (13) | 2.0 | 2.1 (0.9-4.2) |
| Intermediate Wards | 3,890 (55) | 8.5 | 8.6 (3.1-30.7) |
| ICU | 2,326 (33) | 5.1 | 5.4 (1.6-19.2) |
| **Duke Adult Sepsis** | | | |
| ED | 3,509 (38) | 7.7 | 1.3 (0.5-3.1) |
| Intermediate Wards | 3,926 (43) | 8.6 | 15.5 (3.0-51.9) |
| ICU | 1,749 (19) | 3.8 | 4.8 (1.2-24.6) |
| **CDC Adult Sepsis Events** | | | |
| ED | 1,376 (42) | 3.0 | 0.4 (0.2-0.8) |
| Intermediate Wards | 1,282 (39) | 2.8 | 13.1 (2.0-102.0) |
| ICU | 650 (20) | 1.4 | 1.0 (0.1-19.0) |
